# Transmembrane Batten disease proteins interact with a shared network of vesicle sorting proteins to regulate synaptic composition and function

**DOI:** 10.1101/2021.09.16.460691

**Authors:** Mitchell J. Rechtzigel, Brandon L. Meyerink, Hannah Leppert, Tyler B. Johnson, Jacob T. Cain, Gavin Ferrandino, Danielle G. May, Kyle J. Roux, Jon J. Brudvig, Jill M. Weimer

**Affiliations:** Pediatrics and Rare Diseases Group, Sanford Research, Sioux Falls, South Dakota; Department of Pediatrics, Sanford School of Medicine at the University of South Dakota, Vermillion, South Dakota; Basic Biomedical Sciences, Sanford School of Medicine at the University of South Dakota, Vermillion, South Dakota

**Author notes:** co-corresponding authors **Corresponding Author:** Jon J. Brudvig, Sanford Research, 2301 E. 60^th^ N, Sioux Falls, SD 57104, Jill M. Weimer, Sanford Research, 2301 E. 60^th^ N, Sioux Falls, SD 57104. contributed equally (Mitchell J. Rechtzigel, Brandon L. Meyerink).

## Abstract

Batten disease is unique among lysosomal storage disorders for the early and profound manifestation in the central nervous system, but little is known regarding potential neuron-specific roles for the disease-associated proteins. We demonstrate substantial overlap in the protein interactomes of three transmembrane Batten proteins (CLN3, CLN6, and CLN8), and that their absence leads to synaptic depletion of key partners (i.e. SNAREs and tethers) and aberrant synaptic SNARE dynamics *in vivo*, demonstrating a novel shared etiology.

---

Batten disease (also known as Neuronal Ceroid Lipofuscinoses, NCL) is a family of neurodegenerative lysosomal storage disorders caused by mutations in one of at least 13 Ceroid Lipofusinosis Neuronal (CLN) genes^1^. Batten disease is unique among lysosomal storage disorders for the early and profound disease manifestation in the central nervous system, which has frequently been attributed (with little evidence) to a selective vulnerability of neurons to downstream consequences of lysosomal dysfunction. However, while some forms of Batten disease are caused by mutations in genes encoding for lysosomal machinery including catabolic enzymes, other forms are caused by deficiencies in extralysosomal proteins, suggesting that lysosomal dysfunction could be just one consequence of upstream primary defects that are not entirely understood.

In order to gain insights into upstream neuronal functions, we conducted an investigation into the protein interactomes of three transmembrane Batten disease proteins: CLN3, CLN6, and CLN8. Decades of research have uncovered a range of secondary dysfunctions present in cell models of these three disorders, and recent studies in non-neuronal cell models have shed light on their participation in pathways important for lysosomal biogenesis. CLN3 influences the recycling of lysosomal cargo receptors by regulating retromer recruitment and post-Golgi trafficking ^2-4^, and CLN6 and CLN8 cooperatively facilitate the anterograde trafficking of lysosomal cargoes^5,6^. However, reliance on non-neuronal cell models has created potential knowledge gaps in our understanding of neuronal etiology. Synaptic dysfunction precedes lysosomal defects and responds differentially to therapies in some *in vivo* models of these disorders^7-9^, and little is known regarding how these synaptic deficits are manifested.

## CLN3, CLN6, and CLN8 have overlapping protein interactomes enriched for regulators of vesicle identity, fusion, and composition

Given the phenotypic, pathological, and cell biological similarities between CLN3, CLN6, and CLN8 Batten disease, and their overlapping expression patterns in cortical neurons (Fig. 1A), we hypothesized that the three proteins may participate in shared pathways essential for neuronal function. We performed proximity-dependent biotin identification (BioID) screens for each of the three proteins in neuroblastoma cells in order to elucidate their protein interactomes in a neuron-like cell line (Supp. Fig. 1). This identified a large number of enriched proteins in CLN3 (832), CLN6 (515), and CLN8 (1710) samples. Remarkably, there was a substantial degree of overlap between the three groups, with a core set of 266 shared proteins (Fig. 1B). Within this shared interactome, pathway analysis demonstrated an enrichment for proteins involved in vesicle identity, fusion, and composition, (i.e. SNAREs, tethers, and adapter proteins; Figs. 1C and 1D), as well as proteins essential for synaptic vesicle function (Fig. 1E). We confirmed a number of these interactions with co-immunoprecipitations using neuroblastoma cells and mouse cortical lysates, validating the specificity of our BioID screen (Figs. 2A, 2B, and 2C, Supp Table 1).

**Figure 1:**
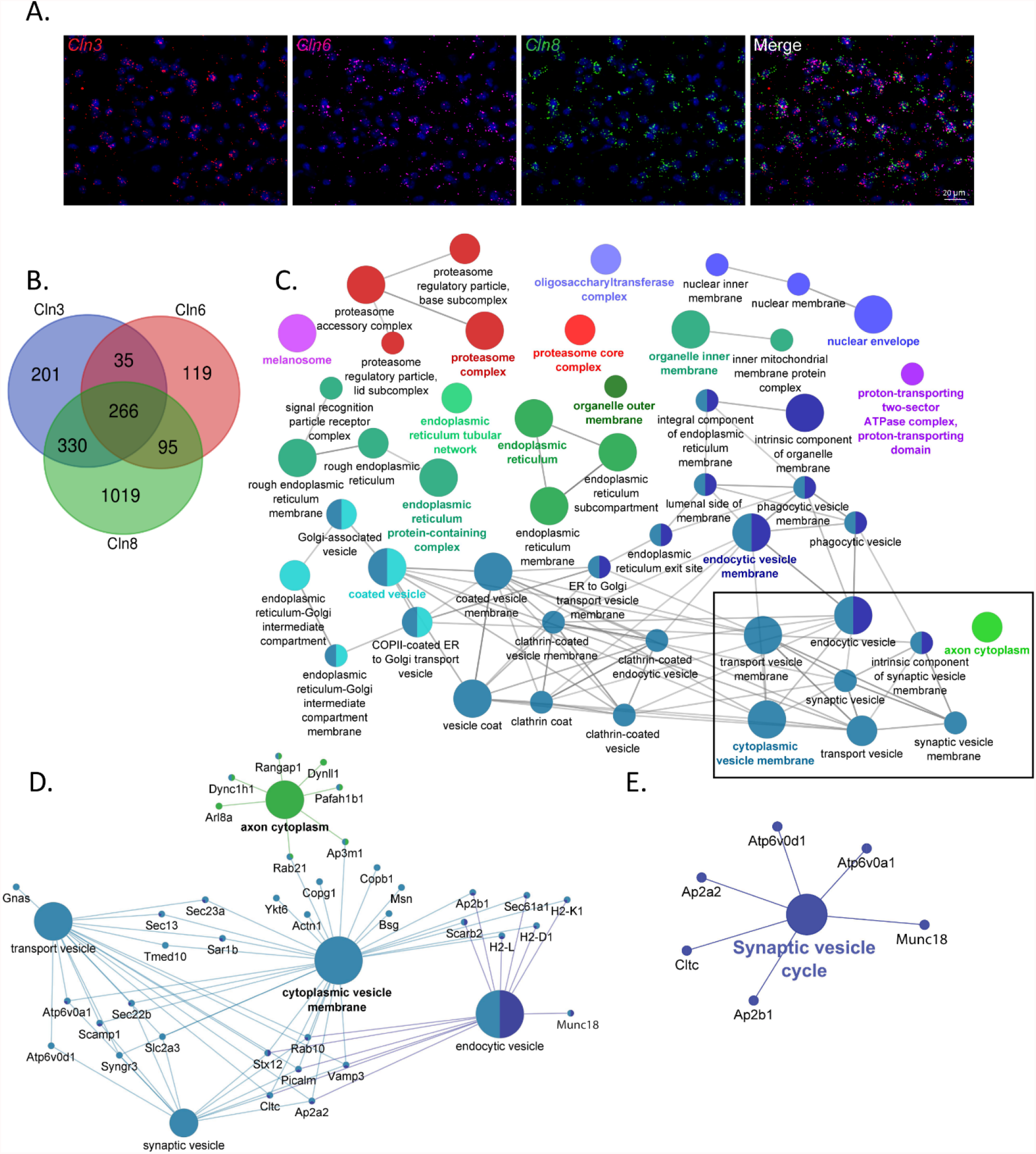
CLN3, CLN6 and CLN8-BioID share a core set of protein interactors with roles in vesicular sorting and presynaptic function. **(A)** RNAscope transcript localization indicates cellular co-occurrence of CLN3, CLN6, and CLN8 in P21 mouse cortex. (B) Venn diagram depicting the number of significantly enriched proteins for each bait protein. (C) ClueGO network of significantly enriched Gene Ontology Consortium (GO) cellular compartment terms of common BioID interactors, with box indicating categories involved in vesicle trafficking and sorting. (D) Shared protein interactors associated with key cellular compartment terms. (E) KEGG Functional terms with corresponding proteins from common BioID interactors.

**Figure 2:**
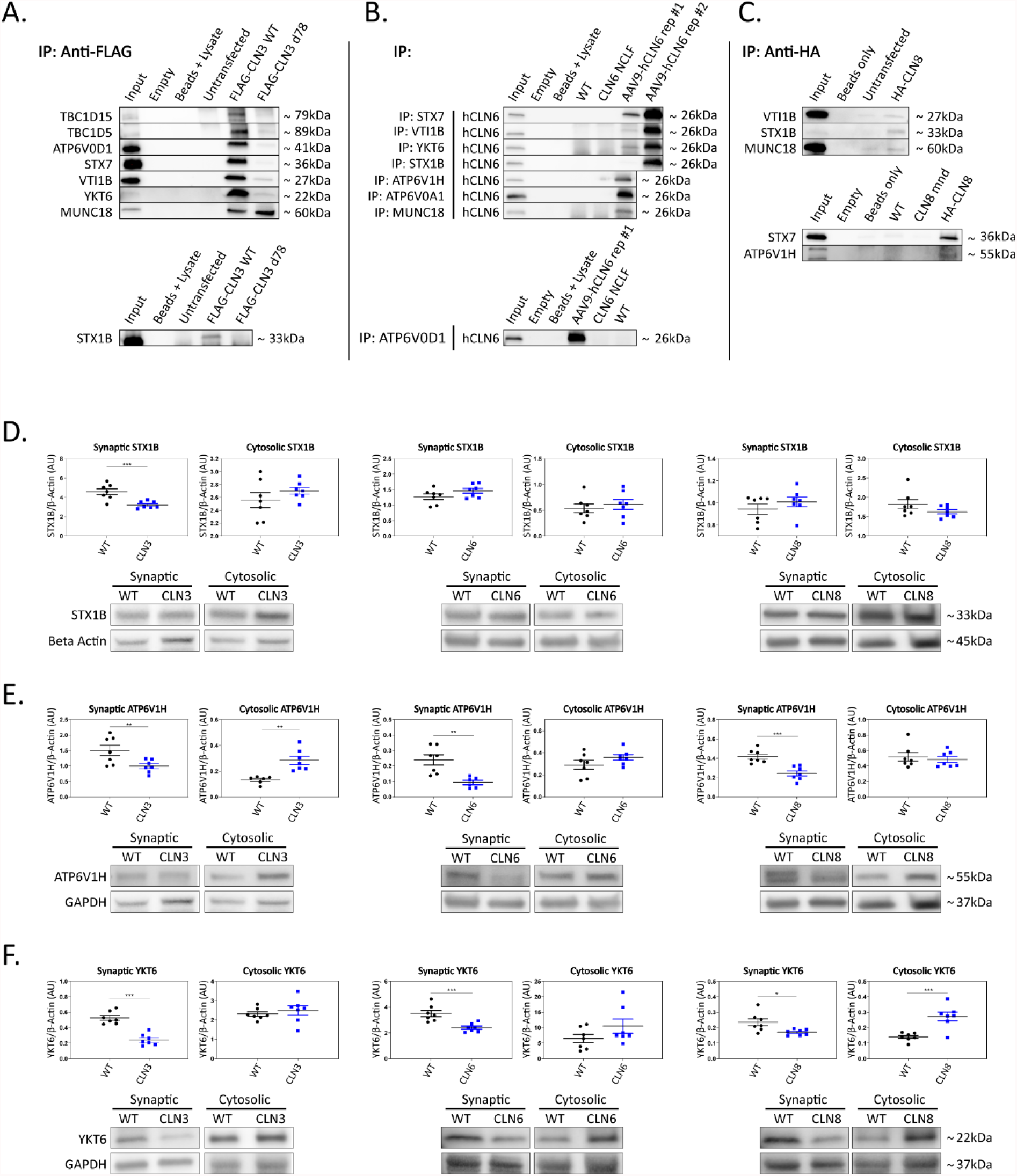
Shared interactors are depleted in cortical synapses. (A) Western blot analyses confirm stable CLN3 interaction with TBC1D15, TBC1D5, ATP6V0D1, STX7, VTI1B, YKT6, MUNC18, and STX1B, (B) stable hCLN6 interaction with STX7, VTI1B, YKT6, STX1B, ATP6V1H, ATP6V0A1, MUNC18, and ATP6V0D1, (C) and stable CLN8 interaction with VTI1B, STX1B, MUNC18, STX7, and ATP6V1H. Due to the lack of commercially available antibodies, tagged CLN3 and CLN8 expression plasmids were used for immunoprecipitation experiments. (D) Analysis of synaptic and cytosolic brain fractions show synaptic depletion of STX1B in *Cln3*^*Δ7/8*^ samples (E), and synaptic depletion of ATP6V1H in *Cln3*^*Δ7/8*^, *Cln6*^*nclf*^, and *Cln8*^*mnd*^ samples coupled with cytosolic accumulation in *Cln3*^*Δ7/8*^. (F) YKT6 was significantly depleted in synaptic fractions of all mutant lines. Also noted was cytosolic accumulation in *Cln8*^*mnd*^ samples. Outliers identified by ROUT analyses, Q=5%. One-tailed t-test (synaptic), two-tailed t-test (cytosolic), *p<0.05, **p<0.01, ***p<0.001, n=7 mice, mean +/- SEM.

## CLN3, CLN6, and CLN8 interactors are depleted in cortical synaptosomes, leading to defects in synaptic vesicle function

To investigate the functional relevance of the shared interactions captured in our screen, we examined the subcellular localization of a subset of proteins with roles in vesicle fusion and presynaptic function. Fresh brain cortices were micro-dissected from postnatal day 30 wild type, *Cln3*^*Δ7/8*^, *Cln6*^*nclf*^, and *Cln8*^*mnd*^ mice (loss of function models for Batten disease^10-12^), followed by synaptic and cytosolic isolation by density separation (Supp Fig. 2F). This revealed striking defects in synaptic composition across all three genotypes (Figs. 2D-2F, Supp Figs. 2A-2E, Supp Table 2). Some targets displayed decreased synaptic abundance in all three genotypes, including ATPase H+ Transporting V1 Subunit H (ATP6V1H, a component of the vacuolar ATPase complex that acidifies lysosomes and synaptic vesicles, facilitating neurotransmitter loading and SNARE-mediated exoctytosis^13-15^; Fig. 2E), Synaptobrevin homolog YKT6 (YKT6, a neuron-enriched SNARE protein localized to a novel vesicular compartment^16^, Fig. 2F), while other targets displayed genotype-specific patterns of depletion, including Syntaxin 7 (STX7, an endosomal Q-SNARE that defines a specialized synaptic vesicle pool in hippocampal neurons^17^, Supp Figs. 2A and 2E) and ATPase H+ Transporting V0 subunit D1 (ATP6V0D1, another component of the vacuolar ATPase that acidifies lysosomes and synaptic vesicles) in *Cln3*^*Δ7/8*^, and mammalian uncoordinated-18 (MUNC18, an essential component of the synaptic vesicle SNARE complex^18^, Supp. Figs. 2D and 2E) in *Cln6*^*nclf*^. In many cases, synaptic depletion was accompanied by cytosolic enrichment, suggesting defects in soma to synaptic terminal sorting or trafficking.

Synaptic terminal composition is tightly regulated and altered stoichiometry of key partners could lead to impaired function. To investigate this possibility, we examined the synaptic vesicle SNARE association state in our disease models. When functioning properly, synaptic vesicles dock and fuse via engagement between vesicular SNAREs on synaptic vesicles (e.g. Synaptobrevin 2, VAMP2) and target snares on the plasma membrane (e.g. Syntaxins 1A and/or 1B, STX1A/STX1B; Synaptosome associated protein 25, SNAP-25). Following neurotransmitter exocytosis, the vesicle fusing ATPase (NSF) dissociates the SNARE complex allowing for vesicle recycling and further rounds of loading and release. Thus, physical associations between these SNARE components can be used to monitor the extent of docking and fusion of synaptic vesicles^19^. We performed SNAP25 coimmunoprecipitations on mouse cortices and quantified levels of bound STX1 and VAMP2, finding prominent aberrations in all three disease models (Figs. 3A-3C). Levels of bound STX1 were significantly increased by at least 2-fold in *Cln3*^*Δ7/8*^, *Cln6*^*nclf*^, and *Cln8*^*mnd*^. VAMP2 trended towards greater levels in *Cln3*^*Δ7/8*^ and *Cln8*^*mnd*^, but was bound at significantly lower levels in *Cln6*^*nclf*^ cortices. These results suggest that synaptic vesicle docking and/or release is compromised in all three of these Batten disease models as a consequence of altered synaptic composition (Fig. 3D).

**Figure 3:**
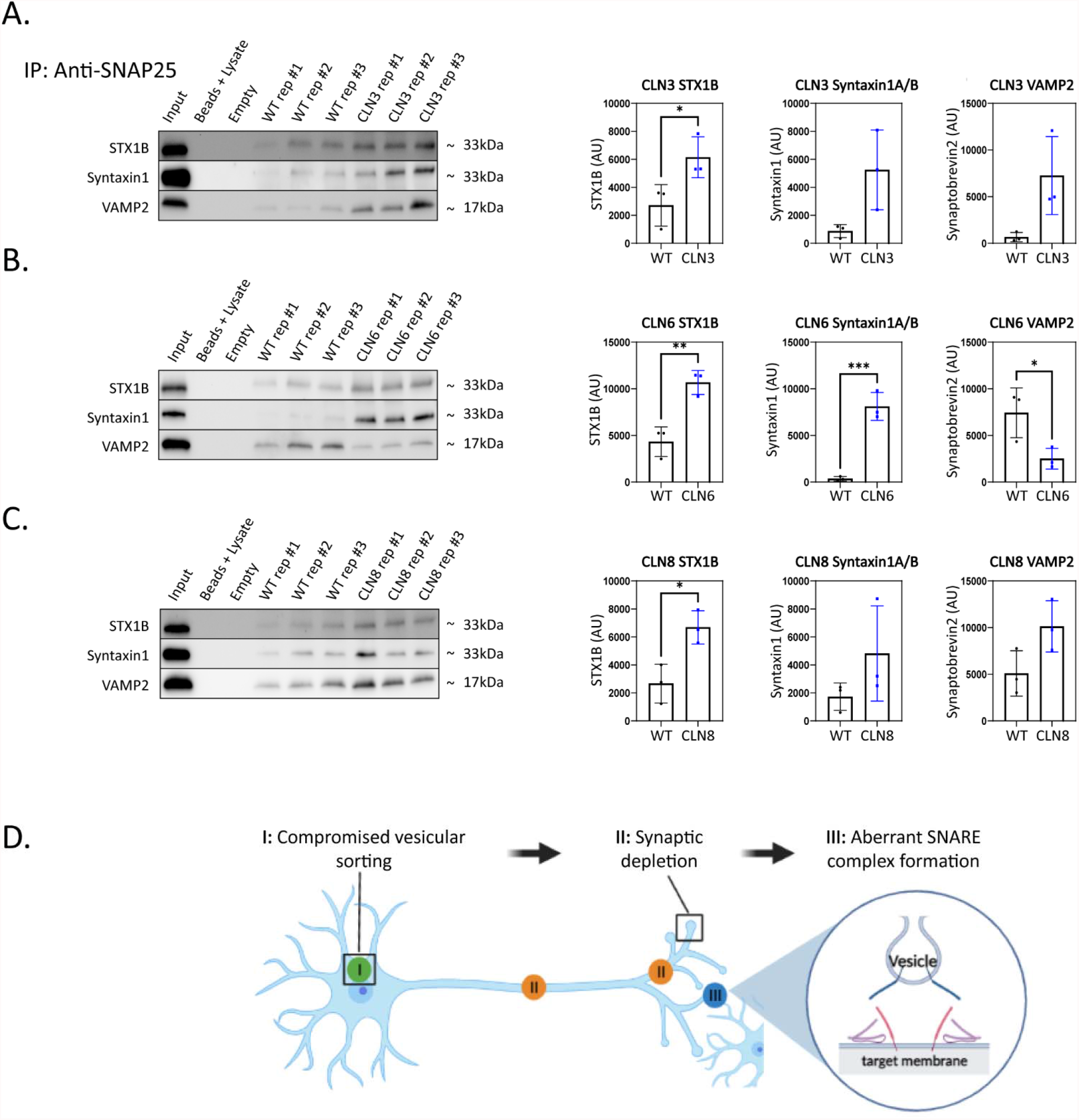
SNAP25 coimmunoprecipitations demonstrate synaptic SNARE dysfunction in *Cln3*^Δ*7/8*^, *Cln6*^*nclf*^, and *Cln8*^*mnd*^ mice. To analyze core SNARE complex formation, P30 mutant brain lysate was immunoprecipitated with anti-SNAP25 antibody and probed for antibodies directed against STX1B, STX1 (recognizes A and B isoforms), and VAMP. (A) Analyses reveal a significant increase in SNAP25-bound STX1B in CLN3, (B) a significant increase in SNAP25 bound STX1B and STX1, and a significant decrease in VAMP2 in Cln6 mutants, (C) and increased binding of SNAP25 and STX1B in CLN8 mutants. (D) The data presented here suggests a model wherein SNARE complex dysfunction is a downstream consequence of (I) comprised vesicular sorting leading to (II) synaptic depletion of targets. (III) In turn, protein imbalance at the synapse results in dysfunctional SNARE complexes formation or release. Two-tailed t-test, *p<0.05, **p<0.01, ***p<0.001, n=3, mean +/- SEM.

Collectively, our results demonstrate a new etiology shared across three neurodegenerative lysosomal storage disorders, and demonstrate novel neuron-specific functions for CLN3, CLN6, and CLN8 in the regulation of synaptic composition and function. The fact that key regulators of vesicle targeting (i.e., SNARES and tethers) are dysregulated in a similar manner to lysosomal proteins (e.g., vacuolar ATPase subunits) provides an intriguing link between seemingly disparate dysfunctions in synapses and lysosomes. Further work will be required to define the molecular-level causes of these defects and the relative contributions the various synaptically depleted interacting proteins. This work suggests that these transmembrane Batten proteins have far-reaching impacts on cellular trafficking and vesicular sorting and supports a multi-faceted disease etiology wherein lysosomal dysfunction is only one salient consequence.

## Online Methods

### Generation of stable expression BioID2 cell lines

Cell lines were maintained in DMEM with 10% fetal bovine serumat 37°C with 5% CO_2_.. Phoenix cells (ATCC #CRL-3213) were transfected with retroviral BioID2^20^ only pBabe, BioID2-KDEL pBabe (translated within the secretory pathway by means of a human albumin signal sequence), BioID2-hCLN3 pRetroX, mCLN6-BioID2 pRetroX, or BioID2-hCLN8 pRetroX using the lipofectamine 3000 lipofection kit (ThermoFisher #L3000008). Viral supernatant was transferred to Neuro2a (ATCC #CCL-131) cells with polybrene (4ug/ml) for 72h transduction prior to selection in maintenance media with puromycin (40 ug/ml) for selection of properly expressing cells.

### Preparation of cell lines and biotinylated samples

BioID protocol was followed as previously described with minor modifications^21^. Briefly, six 10 cm plates were seeded with Neuro2a cells stably expressing BioID2 only, BioID2-KDEL, BioID2-CLN3, CLN6-BioID2, or BioID-CLN8. Control cells received 50⍰μM biotin supplementation for 18h prior to lysis and CLN3/6/8 cells were cultured with doxycycline (20 ug/ml) for 18h prior to supplementation with 50⍰μM biotin for an additional 18h. Cell lysates were pre-cleared with gelatin sepharose 4B (Cytvia, #17095601) prior to overnight pulldown with streptavidin (Cytvia, #90100484). After several washes, biotinylated proteins bound to streptavidin beads were shipped to Sanford-Burnham-Prebys Medical Institute for MS analysis.

### Post-pulldown SDS-Page and Immunoblot

Post-pulldown samples were analyzed via western blot by resuspending 1% of streptavidin beads in SDS-PAGE sample buffer and boiling for 5 min. Proteins were separated on a 4-20% gradient gel (Mini-Protean TGX, Bio-Rad) and transferred to nitrocellulose membrane (Bio-Rad). To detect biotinylated proteins, blots were probed with Streptavidin-HRP (1:5000, ab7403, Abcam) diluted in 0.4% Triton X-100/phosphate-buffered saline and imaged via enhanced chemiluminescent (ECL) substrate detection on LiCor Odyssey FC. Following the detection of biotinylated proteins, HRP activity was quenched via 20-minute incubation in 30% H_2_O_2_. Blots were then blocked with 10% adult bovine serum and 0.2% Triton X-100 in PBS for 30 minutes, and then probed with chicken anti-BioID2 (1:5000, BID2-CP-100, BioFront). An HRP-conjugated anti-chicken antibody (1:10000, ab97135, Abcam) was used to repeat ECL detection.

### Protein Digestion

Biotinylated proteins were digested directly on-beads. Briefly, beads were thawed and cysteine disulfide bonds were reduced with 10 mM tris(2-carboxyethyl)phosphine (TCEP) at 30°C for 60 min followed by cysteine alkylation with 30 mM iodoacetamide (IAA) in the dark at room temperature for 30 min. Following alkylation, urea was diluted to 1 M urea using 50 mM ammonium bicarbonate, and proteins were finally subjected to overnight digestion with mass spec grade Trypsin/Lys-C mix (Promega, Madison, WI). Finally, beads were pulled down and the solution with peptides collected into a new tube. The beads were then washed once with 50mM ammonium bicarbonate to increase peptide recovery. Following digestion, samples were acidified with formic acid (FA) and subsequently desalted using AssayMap C18 cartridges mounted on an Agilent AssayMap BRAVO liquid handling system, C18 cartridges were first conditioned with 100

### Definition of candidate interactors

Candidate interactors for CLN3, CLN6, and CLN8 by the respective BioID2 data set were defined as proteins showing at least 3-fold greater enrichment by label free quantification in CLN-BioID2 samples vs. BioID2 (CLN3 and CLN8) or BioID2-KDEL (CLN6) only samples, in accordance with previous applications of this method^22^.

### Functional analysis of significant interactors

GO analysis and pathway analysis of significant interactors was performed using the DAVID functional annotation tool (https://david.ncifcrf.gov/summary.jsp). Candidate interactors were converted to official gene symbol then selected as the identifier in this tool. Mus musculus was selected as the species of origin. Threshold for the number of genes per term was set to 4 and EASE threshold set to 0.1.

GO network visualization was performed using Cytoscape version 3.8.2 with ClueGO version 2.5.7 and CluePedia version 1.5.7 applications. Significant interactors were input using symbol ID. Organism was set to *Mus musculus*, with grouping based on functional group and fusion of parent and child terms based upon gene abundance. Redundant groups with >50% similar gene identity were merged. Significance defined as p<0.05 with Bonferroni post hoc correction.

### Animals

All animals used for this study were maintained in an AAALAC accredited facility in accordance with IACUC approval (Sanford Research, Sioux Falls, SD). Wildtype (WT), CLN3 (*Cln3*^*Δ7/8*^), CLN6 *(Cln6*^*nclf*^), and CLN8 (*Cln8*^*mnd*^) on C57BL/6J backgrounds were used for *in vivo* and experiments.

### Tissue collection

Mice were sacrificed by carbon dioxide inhalation at a flow rate of 3L/min, diaphragm puncture and cardiac perfusion performed with ice-cold 1X PBS (Corning #21-040-CV). Brains were harvested and cortices collected on dry ice for protein lysates or frozen fresh for RNAscope. For synaptic and cytosolic preparations, the cortices were collected on wet ice.

### RNAscope

*Cln3, Cln6, Cln8, Satb2*, and *Gad67* (ACDBio #497591, 546671, 1029351, 413261, and 413261 respectively) transcript were visualized using ACD RNAscope Multiplex Fluorescent V2 Assay per manufacturers protocol and counterstained with DAPI. Brains were sectioned coronally at 16um with post fixation in 10% Formalin and serial dehydration according to manufacturer’s protocol and mounted with Dako Faramount. Images were captured at 40x on a Nikon Ni-E microscope

### Immunocytochemistry (ICC)

Cells were cultured as outlined above on glass coverslips prior to fixation using 2% PFA/2% Sucrose in 0.1M PBS for 30 minutes. Coverslips were then permeabilized and blocked using blocking solution (0.3% Triton X-100, 1% Bovine Serum Albumin, and 5% goat serum in 0.1M PBS) for 30 minutes followed by incubation with primary antibodies diluted in blocking solution for one hour at room temperature. Coverslips were washed with 0.1M PBS and incubated with secondary antibodies diluted in blocking solution for 30 minutes at room temperature prior to mounting with Dako Faramount media^22^. Cells were imaged using a 60X oil-immersion objective on a Nikon A1 inverted confocal

### Plasmid transfection

Minipreps of cloned plasmids were prepared per manufacturer’s protocol (Zymo Research #D4210). Resulting DNA was transfected into neuroblastoma cell cultures using Lipofectamine transfection following provided protocol (Fisher #L3000015). PCAG-FLAG-CLN3 and pCAG-HA-CLN8 expression plasmids were co-transfected with pCAG-GFP (Addgene #11150) to confirm efficient transfection. Cells were transfected for 24-48 hours, washed 3X with 1X PBS, and collected in cOmplete-Lysis M lysis buffer supplemented with protease and phosphatase inhibitors.

### AAV transduction

scAAV9.CB.CLN6 (AAV9-CLN6) was administered by intracerebroventricular injection into P1 *Cln6*^*nclf*^ animals as previously described. (Cain, 2019)

### Protein Lysis (IP/Western Blotting)

Transfected Neuro2A cells (CLN3/CLN8) and AAV9-CLN6 transduced cortical tissue (CLN6) were homogenized with a pestle homogenizer in 500uL cOmplete Lysis-M lysis buffer (Roche #04719956001) containing protease inhibitors (Roche #04693124001) and phosphatase inhibitors (Sigma #P5726 & P0044). Samples were sonicated on ice for 45 seconds per sample at 30% output (Branson #EDP 100-14-239)

### Brain Fractionation

Cortex samples from P30 WT, *Cln3*^*Δ7/8*^, *Cln6*^*nclf*^, and *Cln8*^*mnd*^ mice were weighed and homogenized in 10mL/gram Syn-PER buffer (Thermo #87793) containing protease and phosphatase inhibitors. Brain fractions were separated by density separation following the manufacturer’s protocol.

### Protein Quantification

Total protein concentration of samples were determined in triplicate using the Pierce BCA Protein Assay Kit (Pierce #23225) following manufacturer’s protocol.

### SDS Page and Immunoblotting

Equal amounts of lysed protein were diluted in water to a total volume of 37.5uL with 1X Laemmli Buffer (BioRad #1610747) supplemented with 10% β-Mercaptoethanol and incubated at 95C for 10 minutes. Samples were loaded on 4-20% Mini-Protean TGX (BioRad #4561096) precast gels and ran at 225V for 30 minutes on ice followed by transfer to nitrocellulose membranes in 20% Methanol Towbin transfer buffer for 60 minutes at 100V. Membranes were washed three times for five minutes in 1X TBS with 0.1% Tween-20, and blocked for 2 hours in TBST supplemented with 5% Adult Bovine Serum. Blots were incubated in primary antibody diluted in 5% ABS blocking buffer overnight at 4C, washed three times, and incubated at 4C for 2 hours in secondary antibody diluted in 5% blocking buffer. Blots were washed an additional three times, exposed to chemiluminescent substrate, and imaged with the ChemiDoc MP imaging system (BioRad #17001402). Antibodies used for immunoblotting described below.

### Immunoprecipitation (CLN3/CLN6)

Due to the lack of a commercially available antibody, FLAG-CLN3-expressing Neuro2A cells were used for immunoprecipitation experiments. 200ug FLAG-CLN3 protein lysate was diluted in a total volume of 500uL with cOmplete-Lysis M lysis buffer (Roche #04719956001) supplemented with protease and phosphatase inhibitors. Protein was incubated with anti-FLAG antibody at 1:50 dilution overnight at 4C.

The following day, samples were added to 25uL pre-washed Pierce A/G magnetic beads (Pierce #88802) and incubated at room temperature for 75 minutes with mixing. Beads were collected using a magnetic stand and flow-through was removed. Beads were washed twice in 1X PBS with 0.05% Tween-20, followed by purified water. Target antigen was eluted at room temperature in 1X Laemmli buffer supplemented with 10% β-Mercaptoethanol for 15 minutes with agitation and analyzed by immunoblotting. CLN6 immunoprecipitation experiments were carried out as described above with AAV9-hCLN6-expressing *Cln6*^*nclf*^ brain lysate. Samples were incubated with target antibody, and immobilized proteins were probed with anti-hCLN6.

### Immunoprecipitation (CLN8)

Neuro2A cell lysate transfected with an HA-tagged CLN8 expression plasmid was used for CLN8 immunoprecipitation experiments. 500uL cOmplete-Lysis M containing 150ug total protein was added to 25uL pre-washed Pierce Anti-HA Magnetic beads (Pierce #88836). Beads were incubated at room temperature for 30 minutes, collected with a magnetic stand, washed, and eluted in 1X Laemmli buffer supplemented with 10% β-Mercaptoethanol. Eluate was analyzed by western blotting protocol described above.

### Antibodies

Antibodies used included anti-ATP6V0A1 (Abnova #H00000535-A01), anti-ATP6V0D1 (Proteintech #18274-1-AP), anti-ATP6V1H (Invitrogen #PA5-22134), anti-Beta Actin (Cell Signaling #4967), anti-BioID2 (BioFront #BID2-CP-100), anti-hCLN6 (proprietary), anti-FLAG (Sigma Aldrich #F1804), anti-GAPDH (Cell Signaling #5174), anti-KDEL (Novus #NBP1-97469), anti-LAMP1 (Abcam #ab24871), anti-MUNC18 (Invitrogen #PA1-742), anti-PSD95 (Cell Signaling #3409), anti-RAB5 (Cell Signaling #3547), anti-SNAP25 (Abcam #ab5666), anti-Streptavidin (Abcam #ab7403), anti-STX1B (R&D Systems #MAB6848), anti-STX7 (Invitrogen #PA5-81945), anti-Synaptobrevin2 (Synaptic Systems #104 211), anti-Syntaxin1 (Synaptic Systems #110 011), anti-TBC1D15 (Proteintech #17252-1-AP), anti-TBC1D5 (Proteintech #17078-1-AP), anti-YKT6 (Abcam #ab236583), anti-Chicken HRP (Abcam #ab97135) anti-Mouse HRP (Cell Signaling #7076S), anti-Rabbit HRP (Invitrogen #31462), anti-Rabbit 568 (Invitrogen #A11011), anti-Mouse 568 (Invitrogen #A21134), anti-Chicken 647 (Invitrogen #A21449).

### Cloning

PCAG-FLAG-CLN3 and pCAG-HA-CLN8 expression vectors were generated using restriction enzyme-based cloning.

### Statistical Analyses

Statistical analyses were performed using GraphPad Prism (v9.1.2; San Diego, CA). Details of the specific tests are noted in the figure legends. In general, one-tailed and two-tailed unpaired t-tests were employed. *p<0.05, **p<0.01, ***p<0.001 Outliers were removed with the ROUT method, Q=5%.

## Acknowledgments

The authors thank Dr. Michelle Hastings for the gift of the FLAG-tagged *Cln3* expression construct. The authors acknowledge the support of the Sanford Research Biochemistry Core (NIGMS CoBRE P20GM106320, and the Histology and Imaging Core (NIGMS CoBRE P20GM103548). Additionally, this work was supported by the National Institutes of Health (NIH # R01NS113233, NIH #R35GM126949) and the ForeBatten Foundation. Additional support by a fellowship to B.L.M. by the USD Neuroscience, Nanotechnology and Networks program through a grant from NSF (DGE-1633213). Figure 3D was created with BioRender.com.

## Author Contributions

The project was conceived by JJB, TBJ, JTC and JMW. JJB, MJR, and BLM designed the experiments. Experiments were performed by MJR, BLM, DM, KJR, HL, and GF. The manuscript was prepared by MJR, BLM, and JJB. JJB, JMW, MJR, BLM, TBJ, JTC HL, GF, and KJR reviewed and edited the manuscript.

---

**Supplemental Figure 1:**
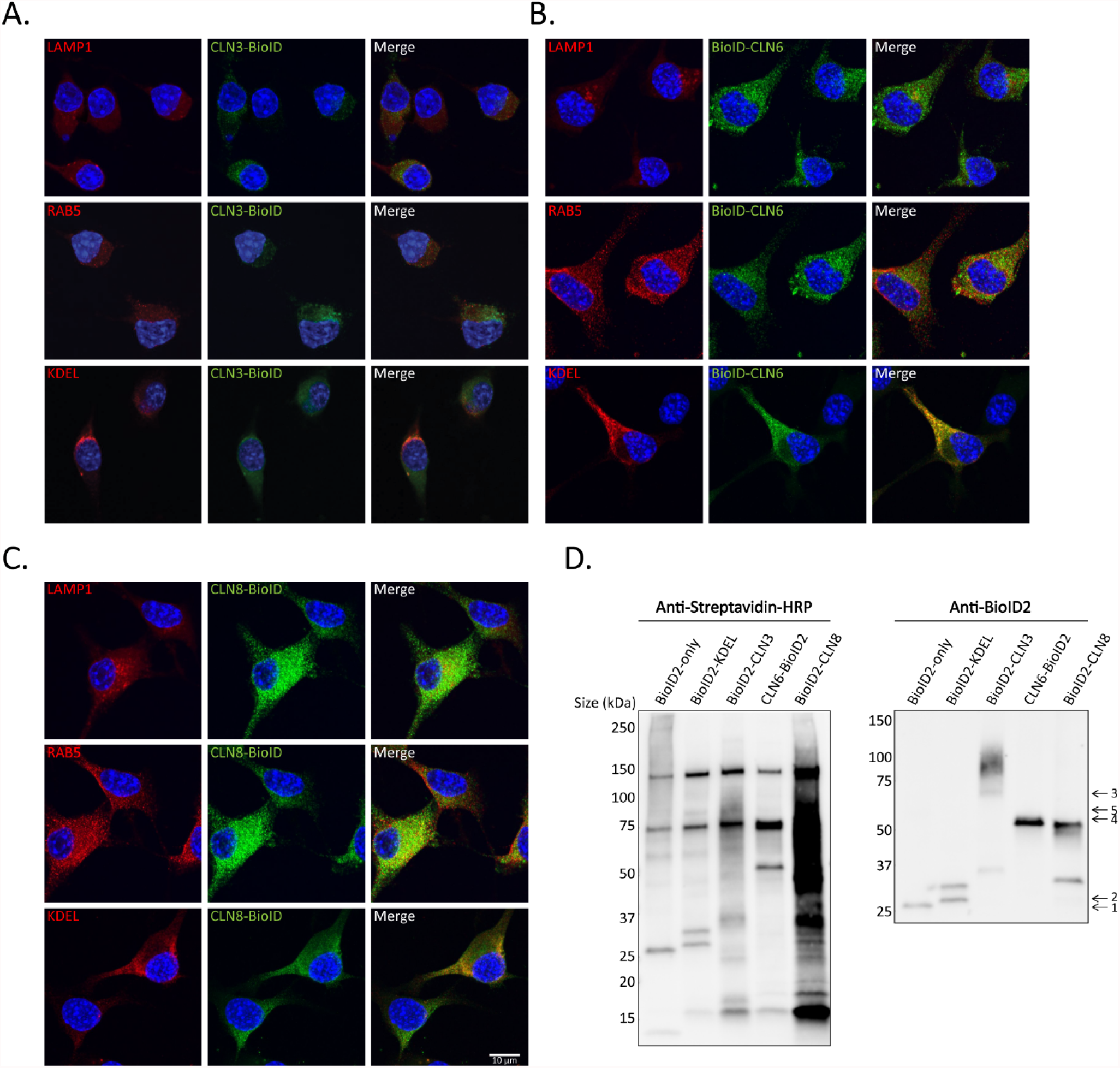
BioID2 expression and protein biotinylation confirmed by IF and western blot. (A) CLN3-BioID, (B) BioID-CLN6, and (C)CLN8-BioID constructs colocalize with markers for lysosomes (anti-LAMP1), endosomes (anti-RAB5) and endoplasmic reticulum (anti-KDEL). Expected band sizes denoted as follows: BioID-only (1) - 26kDa, BioID2-KDEL (2) - 28kDa, BioID2-CLN3 (3) - 68kDa, CLN6-BioID2 (4) - 52kDa, BioID2-CLN8 (5) - 59kDa. Scale bar indicates 10 microns and applies to all images. (D) Western blot analysis of BioID lysates demonstrates successful expression of BioID constructs and resulting biotinylation of proximal proteins.

**Supplemental Figure 2:**
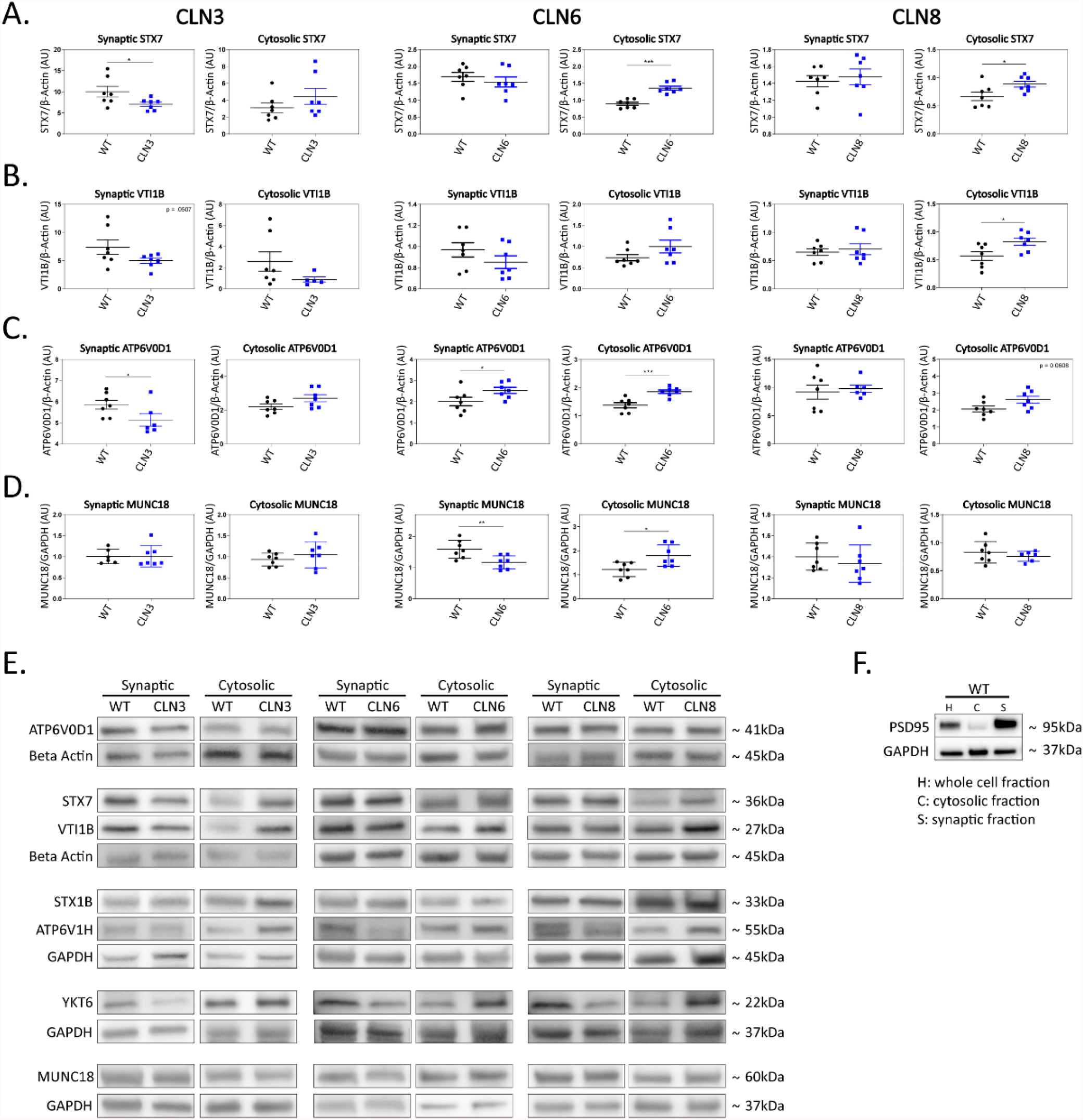
Western blot analysis of P30 mouse cortex brain fractions reveals significant changes in target protein levels when compared to WT. (A) Analysis of target interactor STX7 shows synaptic depletion in CLN3, and cytosolic accumulation in both CLN6 and CLN8. (B) Also demonstrated was cytosolic accumulation of VTI1B in CLN8, (C) synaptic depletion of APT6V0D1 in CLN3 and CLN6, and cytosolic accumulation of ATP6V0D1 in CLN6 mutants. (D) CLN3 and CN8 mutants show no changes in MUNC18 levels, however significant synaptic depletion and cytosolic accumulation was noted in CLN6 mutants. (E) Western blot representative images show visually apparent changes in mutant synaptic and cytosolic fractions compared to WT. (F) Brain fractionation was validated by probing samples for synaptic marker PSD95 and GAPDH to control for protein loading, confirming successful separation of synaptic and cytosolic compartments. Outliers identified by ROUT analyses, Q=5%. One-tailed t-test (synaptic), two-tailed t-test (cytosolic), *p<0.05, **p<0.01, ***p<0.001, n=7 mice, mean +/- SEM.

**Supplementary Table 1:**
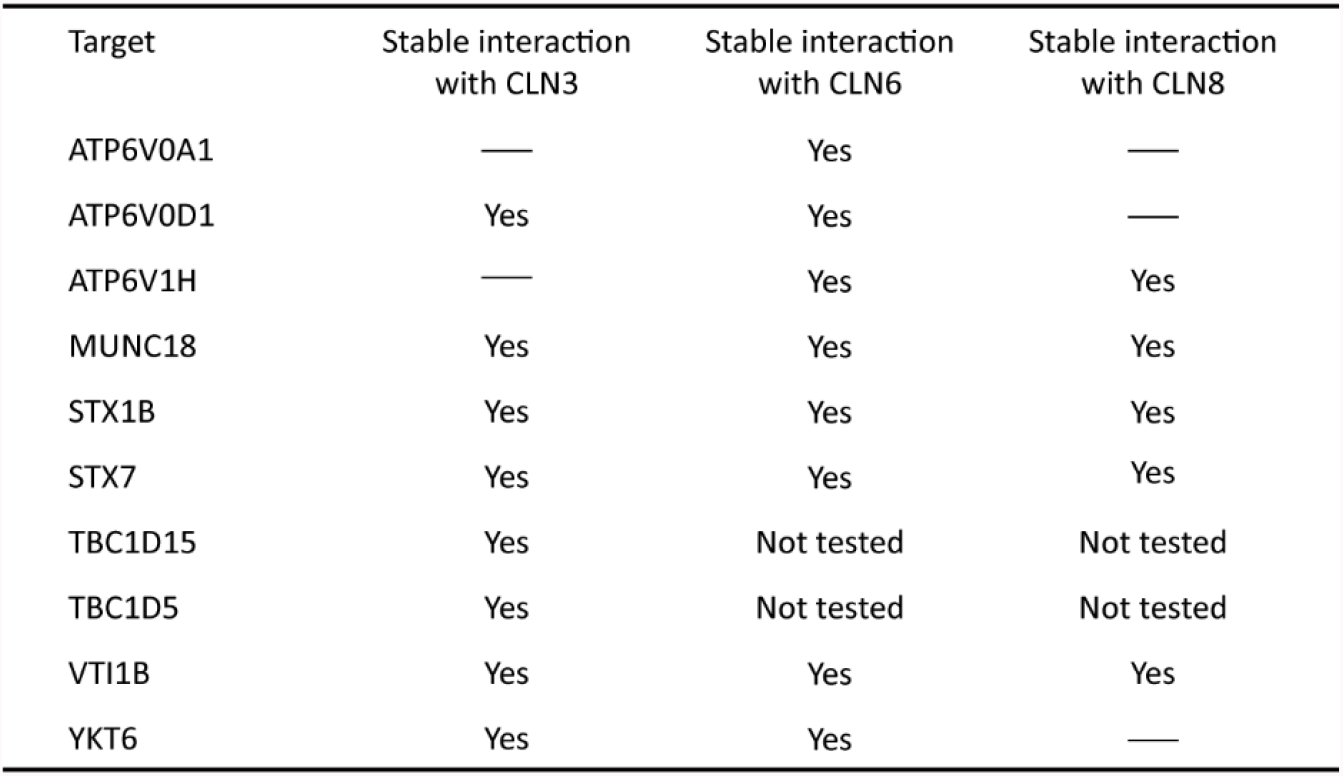
Protein-protein interactions validated by coimmunoprecipitation. Shared BioID-identified interactors of CLN3, CLN6 and CLN8 validated by coimmunoprecipitation. “Yes” denotes stable interaction confirmed while “-” denotes interactions that were not stable under the conditions used for coimmunoprecipitations.

| Target | Stable interaction<br>with CLN3 | Stable interaction<br>with CLN6 | Stable interaction<br>with CLN8 |
| --- | --- | --- | --- |
| ATP6V0A1 | — | Yes | — |
| ATP6V0D1 | Yes | Yes | — |
| ATP6V1H | — | Yes | Yes |
| MUNC18 | Yes | Yes | Yes |
| STX1B | Yes | Yes | Yes |
| STX7 | Yes | Yes | Yes |
| TBC1D15 | Yes | Not tested | Not tested |
| TBC1D5 | Yes | Not tested | Not tested |
| VTI1B | Yes | Yes | Yes |
| YKT6 | Yes | Yes | — |

**Supplementary Table 2:** Synaptic depletion and cytosolic accumulation of targets was observed in P30 mouse cortex brain fractions. Outliers identified by ROUT analyses, Q=5%. One-tailed t-test (synaptic), two-tailed t-test (cytosolic), *p<0.05, **p<0.01, ***p<0.001, n=7, mean +/- SEM

| Supplementary Table 2. Changes in synaptic and cytosolic levels of target proteins in CLN3, CLN6, and CLN8 mutants compared to Wildtype |  |  |  |  |  |  |  |  |  |  |  |  |
| --- | --- | --- | --- | --- | --- | --- | --- | --- | --- | --- | --- | --- |
| Target | CLN3 |  |  |  | CLN6 |  |  |  | CLN8 |  |  |  |
|  | Synaptic | p-value | Cytosolic | p-value | Synaptic | p-value | Cytosolic | p-value | Synaptic | p-value | Cytosolic | p-value |
| ATP6V0D1 | ↓ | .0328 | NS | .0768 | ↑ | .0271 | ↑ | .0009 | NS | .3449 | NS | .0608 |
| ATP6V1H | ↓ | .0089 | ↑ | .0014 | ↓ | .0013 | NS | .1826 | ↓ | .0003 | NS | .6456 |
| MUNC18 | NS | .4750 | NS | .4024 | ↓ | .0043 | ↑ | .0148 | NS | .2210 | NS | .4309 |
| STX1B | ↓ | .0008 | NS | .2792 | NS | .1429 | NS | .5561 | NS | .1584 | NS | .1660 |
| STX7 | ↓ | .0249 | NS | .2470 | NS | .2263 | ↑ | <.0001 | NS | .3335 | ↑ | .0365 |
| VTI1B | ↓ | .0507 | NS | .1616 | NS | .1108 | ↑ | .1436 | NS | .3279 | ↑ | .0269 |
| YKT6 | ↓ | <.0001 | NS | .4962 | ↓ | .0007 | NS | .1554 | ↓ | .0108 | ↑ | .0008 |

## REFERENCES

1 Johnson, T. B. et al. Therapeutic landscape for Batten disease: current treatments and future prospects. Nature reviews. Neurology 15, 161–178, doi:10.1038/s41582-019-0138-8 (2019).

2 Metcalf, D. J., Calvi, A. A., Seaman, M., Mitchison, H. M. & Cutler, D. F. Loss of the Batten disease gene CLN3 prevents exit from the TGN of the mannose 6-phosphate receptor. Traffic (Copenhagen, Denmark) 9, 1905–1914, doi:10.1111/j.1600-0854.2008.00807.x (2008).

3 Yasa, S. et al. CLN3 regulates endosomal function by modulating Rab7A-effector interactions. Journal of cell science 133, doi:10.1242/jcs.234047 (2020).

4 Yasa, S., Sauvageau, E., Modica, G. & Lefrancois, S. CLN5 and CLN3 function as a complex to regulate endolysosome function. The Biochemical journal 478, 2339–2357, doi:10.1042/bcj20210171 (2021).

5 Bajaj, L. et al. A CLN6-CLN8 complex recruits lysosomal enzymes at the ER for Golgi transfer. The Journal of clinical investigation 130, 4118–4132, doi:10.1172/jci130955 (2020).

6 di Ronza, A. et al. CLN8 is an endoplasmic reticulum cargo receptor that regulates lysosome biogenesis. Nature cell biology 20, 1370–1377, doi:10.1038/s41556-018-0228-7 (2018).

7 Ahrens-Nicklas, R. C. et al. Neuronal network dysfunction precedes storage and neurodegeneration in a lysosomal storage disorder. JCI insight 4, doi:10.1172/jci.insight.131961 (2019).

8 Gomez-Giro, G. et al. Synapse alterations precede neuronal damage and storage pathology in a human cerebral organoid model of CLN3-juvenile neuronal ceroid lipofuscinosis. Acta neuropathologica communications 7, 222, doi:10.1186/s40478-019-0871-7 (2019).

9 Ahrens-Nicklas, R. C. et al. Neuronal genetic rescue normalizes brain network dynamics in a lysosomal storage disorder despite persistent storage accumulation. BiorXiv, doi:https://doi.org/10.1101/2021.05.03.442437 (2021).

10 Cotman, S. L. et al. Cln3(Deltaex7/8) knock-in mice with the common JNCL mutation exhibit progressive neurologic disease that begins before birth. Human molecular genetics 11, 2709–2721, doi:10.1093/hmg/11.22.2709 (2002).

11 Morgan, J. P. et al. A murine model of variant late infantile ceroid lipofuscinosis recapitulates behavioral and pathological phenotypes of human disease. PloS one 8, e78694, doi:10.1371/journal.pone.0078694 (2013).

12 Bronson, R. T., Lake, B. D., Cook, S., Taylor, S. & Davisson, M. T. Motor neuron degeneration of mice is a model of neuronal ceroid lipofuscinosis (Batten’s disease). Annals of neurology 33, 381–385, doi:10.1002/ana.410330408 (1993).

13 Bodzeta, A., Kahms, M. & Klingauf, J. The Presynaptic v-ATPase Reversibly Disassembles and Thereby Modulates Exocytosis but Is Not Part of the Fusion Machinery. Cell reports 20, 1348–1359, doi:10.1016/j.celrep.2017.07.040 (2017).

14 Hiesinger, P. R. et al. The v-ATPase V0 subunit a1 is required for a late step in synaptic vesicle exocytosis in Drosophila. Cell 121, 607–620, doi:10.1016/j.cell.2005.03.012 (2005).

15 Poëa-Guyon, S. et al. The V-ATPase membrane domain is a sensor of granular pH that controls the exocytotic machinery. The Journal of cell biology 203, 283–298, doi:10.1083/jcb.201303104 (2013).

16 Hasegawa, H. et al. Mammalian ykt6 is a neuronal SNARE targeted to a specialized compartment by its profilin-like amino terminal domain. Molecular biology of the cell 14, 698–720, doi:10.1091/mbc.e02-09-0556 (2003).

17 Mori, Y., Takenaka, K. I., Fukazawa, Y. & Takamori, S. The endosomal Q-SNARE, Syntaxin 7, defines a rapidly replenishing synaptic vesicle recycling pool in hippocampal neurons. Communications biology 4, 981, doi:10.1038/s42003-021-02512-4 (2021).

18 Verhage, M. et al. Synaptic assembly of the brain in the absence of neurotransmitter secretion. Science (New York, N.Y.) 287, 864–869, doi:10.1126/science.287.5454.864 (2000).

19 Sharma, M., Burré, J. & Südhof, T.C. CSPα promotes SNARE-complex assembly by chaperoning SNAP-25 during synaptic activity. Nature cell biology 13, 30–39, doi:10.1038/ncb2131 (2011).

20 Kim, D. I. et al. An improved smaller biotin ligase for BioID proximity labeling. Molecular biology of the cell 27, 1188–1196, doi:10.1091/mbc.E15-12-0844 (2016).

21 May, D. G. & Roux, K. J. BioID: A Method to Generate a History of Protein Associations. Methods in molecular biology (Clifton, N.J.) 2008, 83–95, doi:10.1007/978-1-4939-9537-0_7 (2019).

22 Brudvig, J. J. et al. MARCKS Is Necessary for Netrin-DCC Signaling and Corpus Callosum Formation. Molecular neurobiology 55, 8388–8402, doi:10.1007/s12035-018-0990-3 (2018).

